# Exploring the role of vascular factors and tissue properties in pulsatile brain deformation

**DOI:** 10.64898/2026.01.23.701278

**Authors:** Marius Burman Ingeberg, Elijah Van Houten, Andrej Shoykhet, Jaco J.M Zwanenburg

**Affiliations:** Translational Neuroimaging Group, Center for Image Sciences, UMC Utrecht, Utrecht, The Netherlands; Université de Strasbourg, CNRS, INSERM, ICube, UMR7357, Strasbourg, France

**Keywords:** Volumetric strain, Octahedral shear strain, Intrinsic Magnetic Resonance Elastography, Strain Tensor Imaging, Human Brain, Brain pulsations, Brain mechanics

## Abstract

**Introduction:** Strain tensor imaging (STI) provides precise measurements of brain tissue deformation caused by cerebral arterial pulsations (CAP). This CAP-related brain tissue deformation is expressed in quantitative strain metrics, such as volumetric strain and octahedral shear strain, which hold promise as quantitative markers of the (mechanical) properties of both the intracerebral vasculature and the intervascular tissue components. However, the extent to which these strain metrics can be specifically linked to the underlying anatomical vascular and tissue properties remains largely unknown. This study aims to explore the relationship between STI metrics and independent markers of pulse pressure (arterial transit time, ATT), vascular function (cerebral blood volume, CBV; cerebral blood flow, CBF; mean transit time, MTT), and tissue properties (shear stiffness).

**Method:** Volumetric and octahedral shear strain were computed from previously obtained 7T displacement data (approximately 2 mm isotropic resolution) of eight healthy subjects (27±7 years). Shear stiffness maps were generated from the same displacement data set using poroviscoelastic intrinsic MR elastography. Regional values of CBV, CBF, MTT, and ATT were obtained from standard-space atlases. Linear mixed-effects models were used to investigate potential regional relationships between specific strain metrics and the corresponding tissue, pulse pressure, or vascular markers.

**Results:** Volumetric strain showed significant positive correlations with CBV (globally, cortical gray and white matter) and significant negative correlations with ATT (globally, and in cortical gray and white matter), but not with shear stiffness. Octahedral shear strain showed a significant negative correlation with shear stiffness (globally, in subcortical gray and white matter) and also with ATT (globally, in cortical gray matter).

**Conclusion:** Volumetric strain reflects mainly vascular properties (pulse pressure, blood volume), while octahedral shear strain is more sensitive to tissue properties. These findings provide a foundation for future studies that investigate the physiological characteristics reflected by these strain metrics and their intricate interplay.

## 1 Introduction

Cerebral arterial pulsations (CAP) driven by the cardiac cycle, generate blood volume changes and subtle tissue deformations throughout the brain. These deformations subsequently drive the movement of cerebrospinal fluid (CSF) (Mestre et al., 2018; Spector et al., 2015; van der Voort et al., 2024) and interstitial fluids (Asgari et al., 2016; Bakker et al., 2016) which are believed to be essential in the clearance of cerebral waste (Tarasoff-Conway et al., 2015). As these pulsatile pressure waves are transmitted through the arterial tree to the level of the microvasculature (Hahn et al., 1996), direct or indirect measurements of pulsatility could probe the microvascular condition, such as in small vessel disease. Additionally, tissue perfusion and viscoelastic metrics have been shown to be highly sensitive to physiological effects, ageing and disease (Hetzer et al., 2018).Thus, methods such as *strain tensor imaging* (STI) (Adams et al., 2020; J. J. Sloots et al., 2021), that quantify strain resulting from such brain tissue deformations using MRI, have the potential to become valuable tools to assess both the vascular and biomechanical state of the human brain.

STI provides, among others, two metrics that may be of high interest in a clinical setting. The first metric is volumetric strain, which describes the relative expansion or compression of a voxel during CAPs. The second metric is octahedral shear strain, which reflects the deformation of a voxel independent of expansion or compression. These metrics are of particular interest as they reflect the state of- and interaction between-vascular and intervascular tissue components of the brain. Meanwhile, the measurement of these strain metrics remains a relatively recent technical achievement, and their specificity to the underlying biological and physiological drivers are still largely unknown. Given the brain’s highly dynamic and complex nature, these drivers are likely multifactorial and interdependent. Several factors may contribute to the strain observed in a given voxel, including blood pressure gradients, blood volume fraction, brain tissue stiffness, and vascular wall compliance, among others. Better understanding of the relationship between the strain metrics and the underlying drivers is important for advancing our knowledge of both normal brain function and pathophysiology, particularly in relation to diseases that affect the microvasculature. This, in turn, is an essential first step towards utilizing these metrics in clinical research.

Recent work has achieved high-quality brain tissue strain measurements by leveraging 7T imaging with Displacement Encoding with Stimulated Echoes (DENSE) (Aletras et al., 1999). This sequence achieves high signal-to-noise ratio (SNR) (Adams et al., 2020) because its displacement encoding is optimized for submillimeter motions - the same order of magnitude as the tissue displacements produced by brain pulsations. Its sensitivity to displacements of the same magnitude as brain pulsations gives DENSE an advantage over other phase contrast techniques. With DENSE, Adams et al. (2020) observed a 2.3-fold difference in peak volumetric strain between gray and white matter, though the underlying cause remains unclear. It was speculated that the differences between tissue types likely result from variations in both blood volume and tissue viscoelasticity. Since then, intrinsic MR elastography (iMRE) has been shown to be feasible, enabling the estimation of shear stiffness from the same displacement measurements used to assess brain tissue strain (Burman Ingeberg et al., 2023, 2025). Shear stiffness has previously been shown to be a reliable indicator of the mechanical integrity of neuronal tissue (Hirsch et al., 2013; Wuerfel et al., 2010), positioning iMRE to potentially disentangle vascular contributions from tissue properties and provide deeper insight into the mechanisms underlying brain strain.

This study aims to investigate potential drivers contributing to volumetric strain and octahedral shear strain during CAPs in the human brain, with a particular focus on distinguishing vascular influences from those related to tissue stiffness. We considered three complementary categories of drivers: (1) pulse pressure effects, approximated by arterial transit time (ATT); (2) vascular and hemodynamic factors, captured by CBV, cerebral blood flow (CBF), and mean transit time (MTT); and (3) tissue properties, represented by shear stiffness. A regional analysis across gray matter (subdivided into cortical and subcortical) and white matter was performed to examine how these drivers influence strain. This study serves as a foundational exploration of drivers of strain, providing a baseline for more focused and specific future experiments.

## 2 Method

### 2.1 Acquisition

The analysis presented here is based on displacement data previously collected by Adams et al. (Adams et al., 2020). A detailed description of the data acquisition method is available in the original publication; only a brief summary is provided here. All participants provided written informed consent, and the study was approved by the institution’s ethical review board.

Displacement data were acquired from eight healthy young adults (three females; mean age: 27 ± 6 years) using a Displacement Encoding with Stimulated Echoes (DENSE) sequence on a 7T MR scanner (Philips Healthcare) with a 3D EPI readout scheme. Repeated scans were performed, with subjects briefly exiting the scanner for up to 10 minutes between sessions. Retrospective gating synchronized the DENSE measurements with the cardiac cycle, which was detected using pulse oximetry. The spatial resolution of the acquired images was 1.95 mm × 1.95 mm × 2.2 mm (FH x AP x RL), and 20 temporal phases were reconstructed per cardiac cycle. Three separate acquisitions were performed with displacement encoding (*D*_*enc*_) sensitivities of 0.175 mm, 0.175 mm, and 0.35 mm in the anterior-posterior (AP), right-left (RL), and foot-head (FH) directions, respectively. The acquisition of each motion encoding direction took 2:24 minutes, for an average heart rate of 60 beats per minute. Additionally, T1-weighted images with a resolution of 0.93 mm × 0.93 mm × 1.0 mm were acquired during both scanning sessions to facilitate segmentation and registration. Tissue probability maps were generated from the T1-weighted images using the Computational Anatomy Toolbox (Jena University Hospital, Departments of Psychiatry and Neurology) extension for SPM12 (Wellcome Trust Centre for Neuroimaging, University College London).

Because of the possibility of subject motion between the three DENSE acquisitions, images from the different motion encodings were registered to the same space. This was done by first rigidly registering the AP and FH magnitude images to the RL magnitude images. Then, the last RL image of the cardiac cycle was non-linearly registered to the corresponding T1-weighted image for each subject. Finally, all images were transformed to the T1-weighted space using the transformation parameters given by the transformation from the RL magnitude image to the T1-weighted image.

Motion artifacts related to physiological noise were identified following the methodology described by Adams et al. (Adams et al., 2020). In short, SNR maps were created by scaling DENSE magnitude images from the end of the cardiac cycle (containing the majority of artifacts) by the temporal standard deviation. The SNR maps were then thresholded to create artifact masks, which were applied to the DENSE measurements. The repeated scan of Subject 7 was excluded as it contained more than twice the number of voxels with motion artifacts in the displacement measurements compared to the average of all other scans.

### 2.2 Strain computation

The DENSE images were converted to displacement fields by scaling the phase images with the corresponding *D*_*enc*_ value for each displacement encoding direction. Displacement gradient fields were then computed by taking the spatial derivatives of the AP, RL, and FH displacement maps along their respective directions. Phase wraps that occurred due to large numeric derivatives were unwrapped under the small-strain assumption, which assumes that displacement differences between neighboring voxels are smaller than the tag spacing. Hence, the maximum tolerated strain was

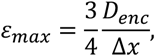

with Δ*x* being the distance between neighboring voxels (J. J. Sloots et al., 2021), which for the used data is 6.7%, 6.0% and 13% for AP, RL and FH directions, respectively. The deformation gradient tenor **F** was then computed as **F** = ∇**u** + **I**, where ***u***_***x***_, ***u***_***y***_, and ***u***_***z***_ are the displacement fields in the RL, AP and FH directions respectively, and **I** is the identity matrix. Finally, the Lagrangian strain tensor **E** was constructed as

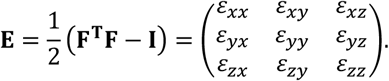

Two properties of interest can be derived from the tensor **E**, the volumetric strain and the octahedral shear strain. Volumetric strain reflects the net expansion or compression of a given voxel, and is calculated from the diagonal of **E** as

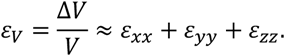

Octahedral shear strain reflects the deformation of a voxel, independent of the volumetric strain, and is calculated as (McGarry et al., 2011)

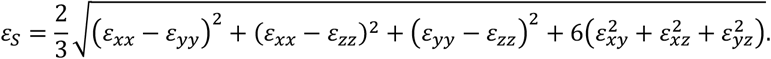

Zero octahedral shear strain indicates isotropic expansion or compression of a voxel, while larger values indicate shear deformation. Both volumetric and octahedral shear strain are independent of the orientation of the coordinate system (McGarry et al., 2011). Absolute volumetric strain values exceeding 2% and octahedral shear strain values exceeding 4% were attributed to partial-volume effects with CSF, large blood vessels, or residual artifacts and were excluded from analysis (Adams et al., 2020). As the magnitude of these strains vary over the cardiac cycle, peak strains were chosen as the metric of interest.

### 2.3 Tissue stiffness estimation

Brain tissue stiffness was estimated using poroviscoelastic inversion for iMRE. This method had been applied in a prior analysis, with full details provided in the original publications (Burman Ingeberg et al., 2025). A brief summary of the approach is presented here.

The displacement fields of each subject were transformed into frequency space by applying the Fast Fourier Transform to the temporal phases for each voxel. The motion at the first harmonic frequency, which roughly corresponds to 1 Hz (for a cardiac rhythm of 60 bpm), was selected for estimation of shear stiffness. Tetrahedral finite element models were created from the corresponding displacement maps for each subject. The finite element models were then used in a subzone-based nonlinear inversion scheme (McGarry et al., 2012; Van Houten et al., 2001) to estimate the underlying mechanical properties using a poroviscoelastic model. With such a model, the brain is modeled as a biphasic material that comprises a viscoelastic porous matrix which is permeated by a viscous pore fluid. A total of 600 global iterations were allowed for property map of each subject to ensure convergence. The shear stiffness was calculated from the recovered complex shear modulus, *G*^∗^, as

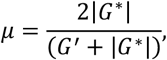

where *G*^′^ is the real part of *G*^∗^.

### 2.4 Comparison with ATLAS maps

The dependence of volumetric strain and octahedral shear strain on vascular and pulse-pressure-related properties was investigated by comparison to standardized atlas maps in MNI space reported in literature. The following maps were used for analysis: a cerebral blood volume (CBV) map that was generated using 134 non-linearly registered CBV maps obtained from DSC MRI (Fuster-Garcia et al., 2021); cerebral blood flow (CBF) and arterial transit time (ATT) maps obtained using high-resolution whole brain arterial spin labeling (ASL) for 10 volunteers (Taso et al., 2021); a mean transit time (MTT) map, distributed as part of the *rapidtide* analysis package (Frederick B. Blaise, 2016). This final map was derived using the procedure described in the poster Blaise et al., 2017, updated to use the data from 4036 resting state scans from the HCP1200 release. A detailed overview of sources and subject characteristics can be found in Table S1 in supplementary files. CBF and MTT are both convoluted metrics, relating to multiple underlying factors (such as blood volume, arterial pressure and hematocrit), which in turn hampers interpretation. In contrast, CBV corresponds to a more unambiguous representation of vascular anatomy. For this reason, we focus on CBV in the main analysis, with CBF and MTT presented in the supplementary material for completeness. Each map was registered to each respective subject using Elastix (Klein et al., 2010), with separate registrations for the repeated scans. This was done by first rigidly registering the ICBM2009c non-linear average T1-weighted template to the T1-weighted image of each respective subject, followed by an affine and non-linear registration with nearest neighbor interpolation. Figure 1 shows all properties of interest for a representative subject.

**Figure 1.**
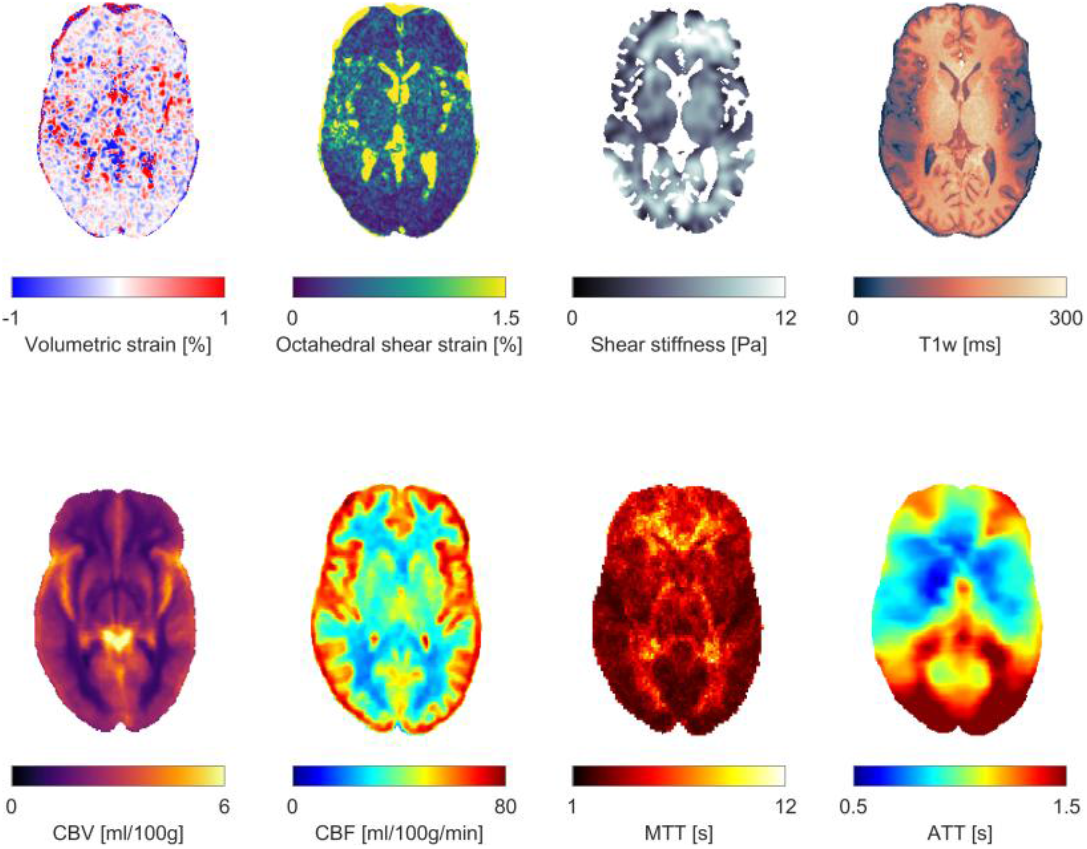
Representative axial slices of the properties of interest along with a T1-weighted image for anatomical reference for one subject. The top row contains subject specific measurements, while the bottom row contains standardized mean atlas maps that have been registered to subject space. Volumetric and octahedral shear strain maps are shown at a timepoint corresponding to peak strain.

### 2.5 Regional extraction

The potential dependence of volumetric strain and octahedral shear strain on microstructure and vascular properties was investigated by performing a regional analysis. A total of 28 regions were investigated, categorized based on the type of tissue: cortical gray matter (GM), subcortical GM, and white matter (WM). Cortical and subcortical regions were identified using the CerebrA atlas (Manera et al., 2020), and WM regions were identified using the JHU-ICBM-tracts and JHU-ICBM-labels 1 mm atlases (Mori et al., 2008; Mori S et al., 2006). Table 1 contains all regions used in this analysis, with their respective abbreviation, and tissue type. Partial volume effects were minimized by eroding each region by 1 voxel.

**Table 1.**
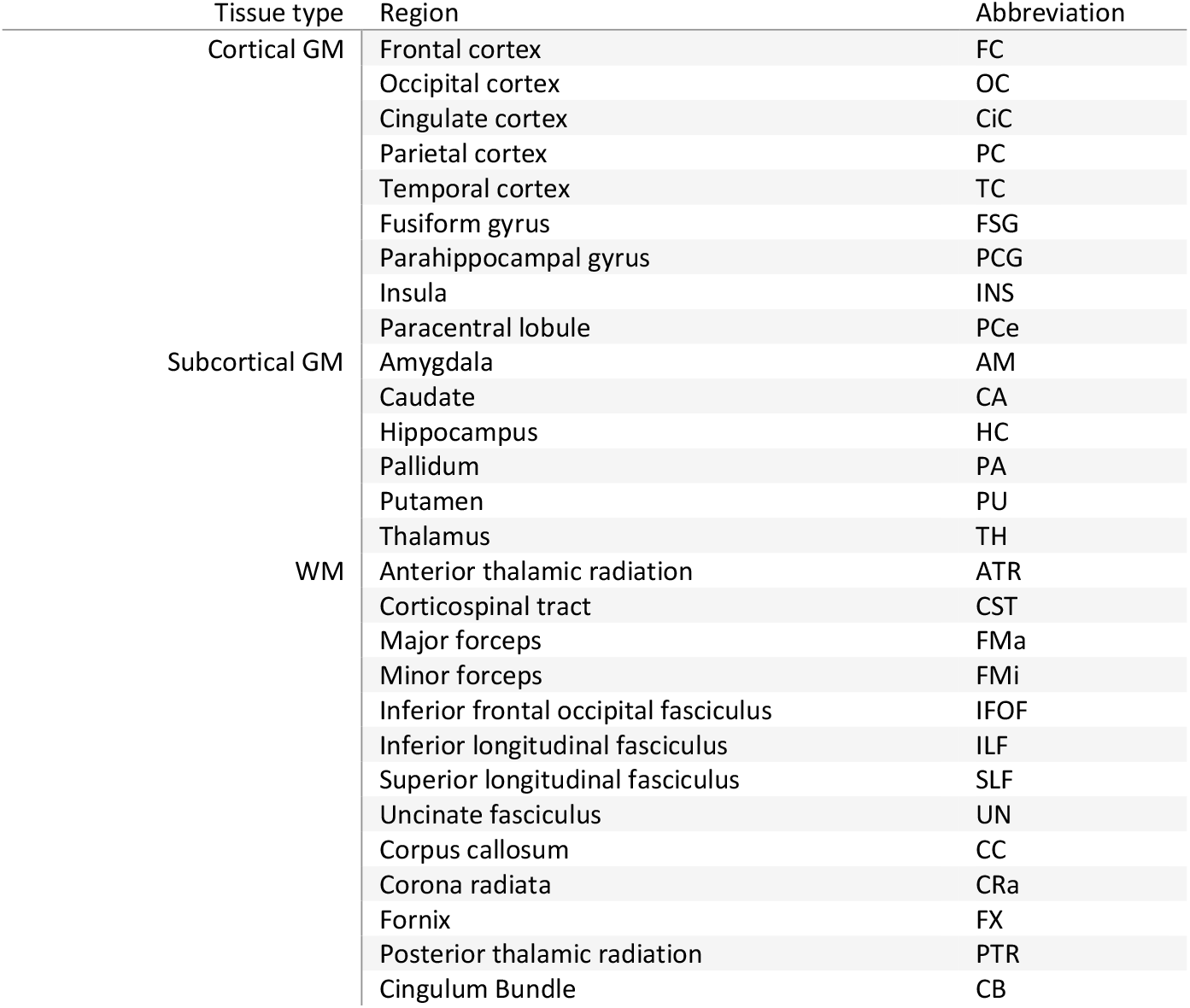
All regions of interest and their respective abbreviations, categorized into tissue types.

Voxels with less than 95% probability of being classified as either GM or WM were excluded from the analysis to avoid CSF and partial-volume effects at tissue boundaries. To ensure mutually comparable regional correlations across the various properties, only voxels for which all parameters were available were included. Voxels corresponding to missing parameters, arising from the removal of artifactual strains in the strain maps or from problematic fluid flow in the shear stiffness map (Burman Ingeberg et al., 2023) (see Figure 1), were excluded from all property maps to maintain consistency in regional comparisons. Consequently, the number of voxels per region was considerably reduced, particularly in gray matter areas near CSF. Therefore, regions with fewer than an average of 300 voxels across subjects or those where more than 80% of voxels were removed (compared to the eroded region mask) were excluded from the analysis. The mean value of all properties of interest was computed for each region. For time-dependent properties such as volumetric strain and octahedral shear strain, the mean regional value was computed for each time point. Peak-to-peak volumetric strain *ε*_*V*,pp_ and octahedral shear strain *ε*_*S*,pp_ were then computed by taking the difference between the maximum and minimum value across time.

### 2.6 Statistical analysis

The regional relationships between the measured strains and the vascular and tissue properties were investigated using linear mixed effects models. This was done separately for cortical GM, subcortical GM and WM, as well as globally (all regions). Each subject was specified as a random effect, with repeated measurements being specified as a nested random effect within each subject. The model was weighted by the size of each region, ensuring that regions with larger volumes had a greater influence on the estimation. Data points with an absolute z-score over three were considered outliers and were removed from the analysis. Bonferroni correction was used to correct for multiple comparisons such that statistically significant effects were determined at 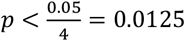.

## 3 Results

All regions except the Fornix, the Uncinate fasciculus, the Pallidum, and the Temporal cortex met the requirements to be kept for analysis for all subjects. The mean regional values for each property of interest are presented in Table 2, averaged across measurements.

**Table 2.**
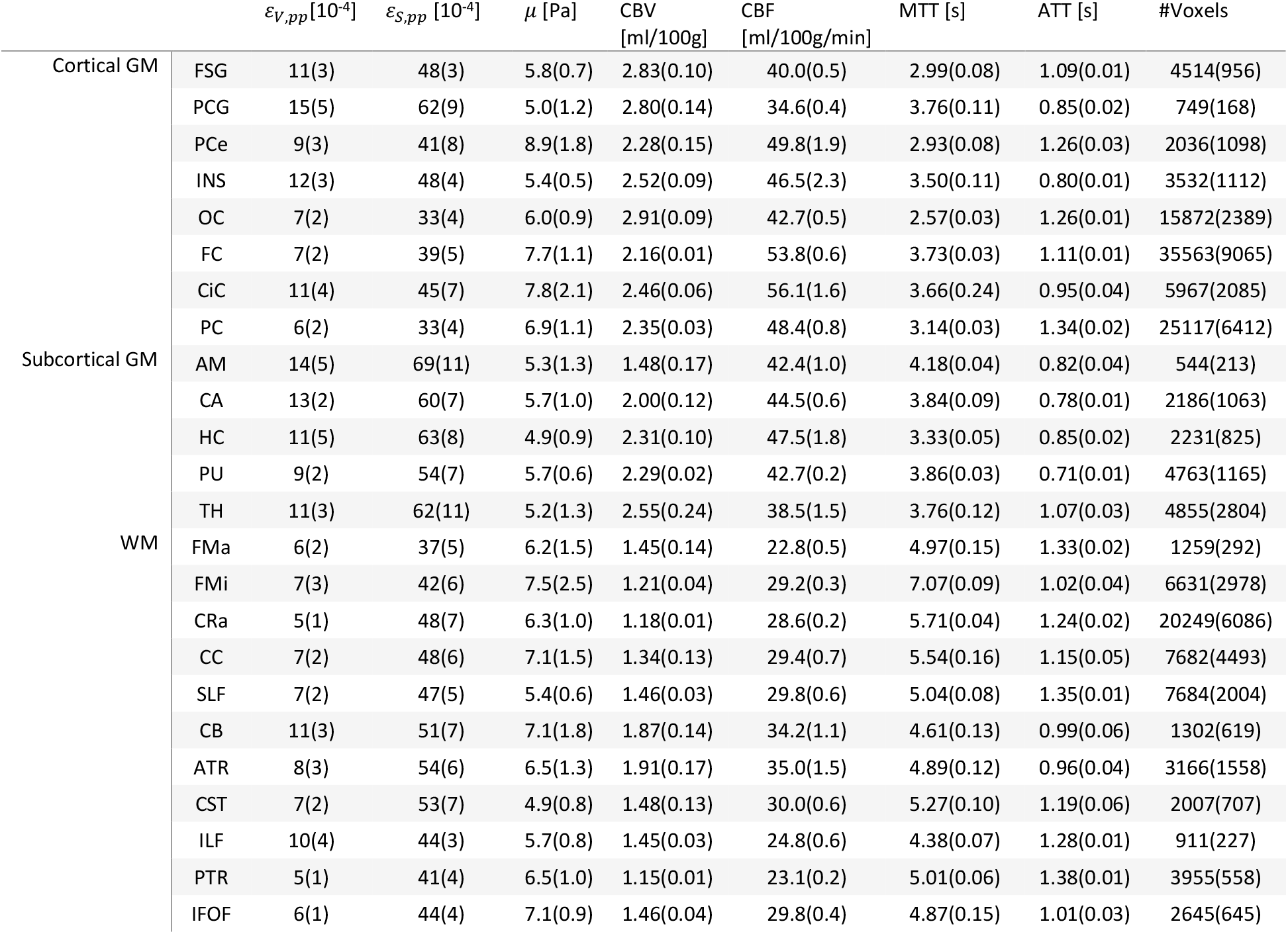
Mean and standard deviation of regional values across all measurements for all properties, along with the mean number of voxels of each region. The standard deviations of atlas properties reflect variations caused by registration differences and the variable exclusion of voxels across subjects.

### 3.1 Volumetric strain

#### 3.1.1 Relationships with ATT (proxy for pulse pressure)

The relationships between volumetric strain and ATT are illustrated in Figure 2, with each data point representing the subject-wise mean regional values. The corresponding regression statistics for each tissue type are detailed in Table 3. Outlier analysis excluded two regional values. Volumetric strain demonstrated a significant negative trend in cortical gray matter, white matter and at the global level.

**Table 3.** The intercept and slope (both with 95% confidence intervals), *p-*value, and *R*^2^ value obtained from linear regressions between volumetric strain and ATT using a linear mixed effects model for each tissue type. Significant correlations after Bonferroni correction are indicated in bold.

| Region | Intercept [$10^{-4}$] | Slope [$10^{-4}$] | $p$-value | $R^2$ |
| --- | --- | --- | --- | --- |
| Global | 16.7 (9.1, 14.8) | -8.1 (-7.1, -2.5) | **< 0.001** | 0.43 |
| Cortical GM | 18.4 (15.3, 21.6) | -9.3 (-11.8, -6.9) | **< 0.001** | 0.59 |
| Subcortical GM | 11.6 (7.7, 15.5) | -1.4 (-5.7, 2.9) | 0.521 | 0.23 |
| WM | 12.0 (9.1, 14.8) | -4.8 (-7.1, -2.5) | **< 0.001** | 0.29 |

**Figure 2.**
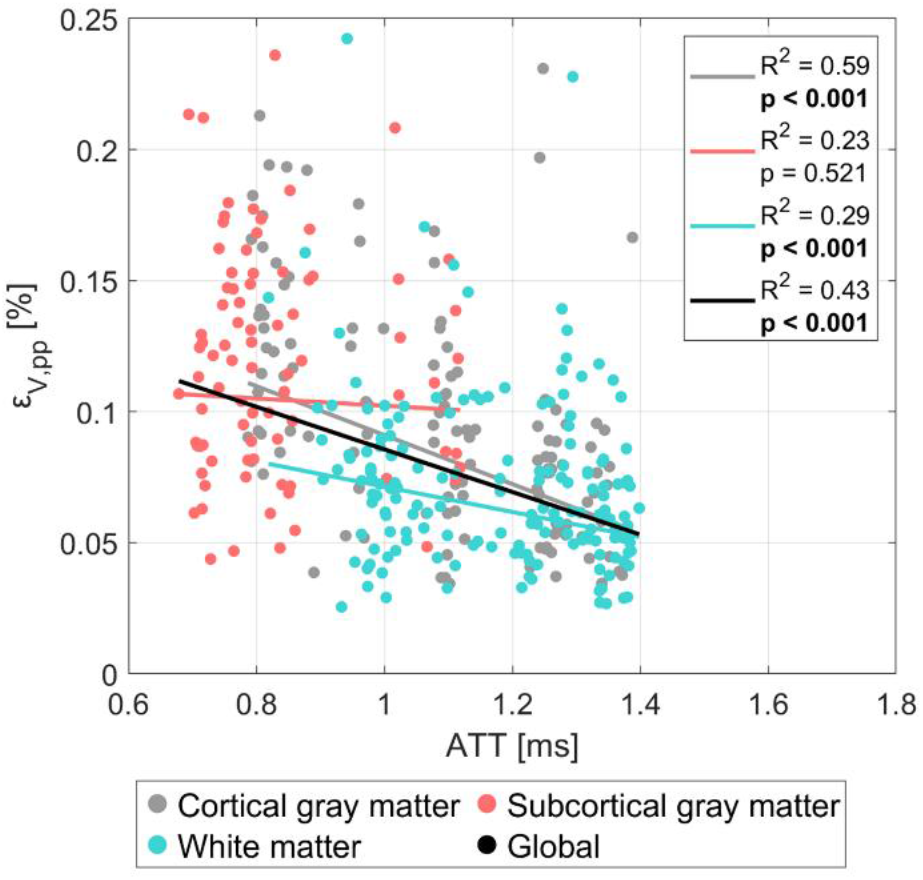
The dependence of volumetric strain on ATT for cortical gray matter (gray), subcortical gray matter (red), white matter (teal). Each dot represents the subject-wise mean value across a given region, calculated only for one repeated scan. For each tissue type, the colored line shows the fixed-effect linear regression (weighted by region size) estimated from a linear mixed effects model, representing the overall population-level trend while accounting for subject- and scan-specific variability. The black line depicts the equivalent regression across all tissue types combined. The legend displays the corresponding *p*-values and *R*^2^ for each regression, where significant correlations after Bonferroni correction are indicated in bold.

#### 3.1.2 Relationships with CBV (proxy for vascular structure)

The relationships between volumetric strain and CBV are illustrated in Figure 3, with each data point representing the subject-wise mean regional values. The corresponding regression statistics for each tissue type are detailed in Table 4. A separate outlier analysis was performed for the CBV comparison, with three regional values excluded. Linear regression analysis revealed significant positive correlations between volumetric strain and CBV in WM, cortical GM and globally. However, a significant negative trend was observed in subcortical GM, potentially indicating distinct tissue-specific dependencies.

**Table 4.** The intercept and slope (both with 95% confidence intervals), *p-*value, and *R*^2^ value obtained from linear regressions between volumetric strain and CBV using a linear mixed effects model for each tissue type. Significant correlations after Bonferroni correction are indicated in bold.

| Region | Intercept [$10^{-4}$] | Slope [$10^{-4}$] | $p$-value | $R^2$ |
| --- | --- | --- | --- | --- |
| Global | 3.9 (-1.0, 2.8) | 1.7 (2.7, 5.3) | **< 0.001** | 0.33 |
| Cortical GM | 1.8 (-2.0, 5.5) | 2.4 (0.9, 3.8) | **0.002** | 0.40 |
| Subcortical GM | 18.4 (14.1, 22.7) | -3.4 (-5.2, -1.7) | **< 0.001** | 0.42 |
| WM | 0.9 (-1.0, 2.8) | 4.0 (2.7, 5.3) | **< 0.001** | 0.37 |

**Figure 3.**
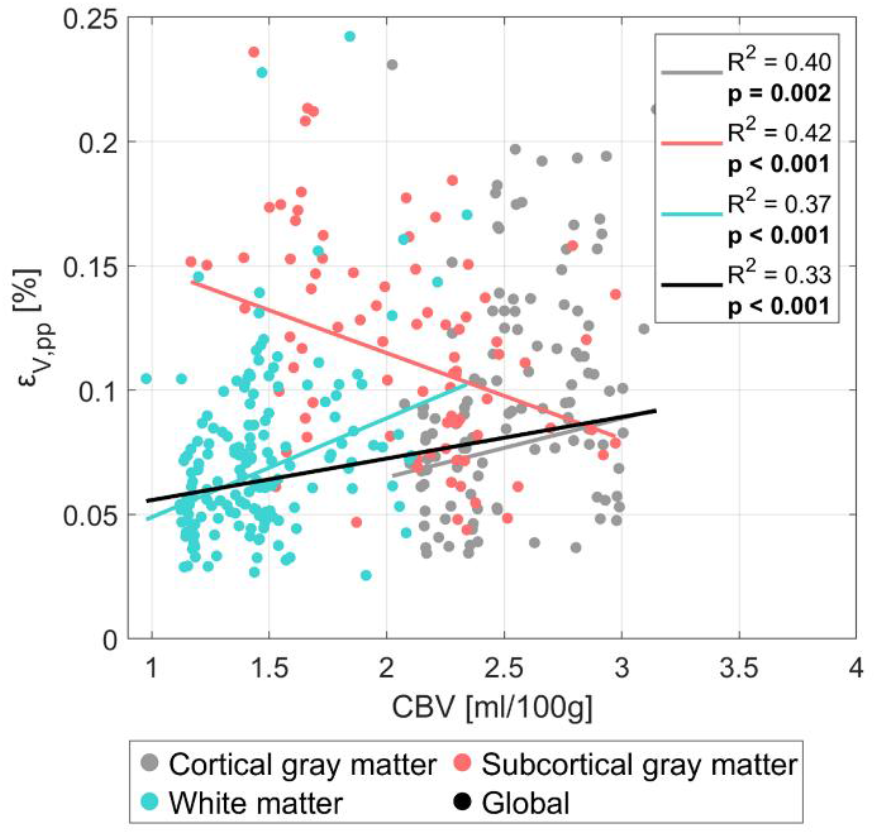
The dependence of volumetric strain on CBV for cortical gray matter (gray), subcortical gray matter (red), white matter (teal). Each dot represents the subject-wise mean value across a given region, calculated only for one repeated scan. For each tissue type, the colored line shows the fixed-effect linear regression (weighted by region size) estimated from a linear mixed effects model, representing the overall population-level trend while accounting for subject- and scan-specific variability. The black line depicts the equivalent regression across all tissue types combined. The legend displays the corresponding *p*-values and *R*^2^ for each regression, where significant correlations after Bonferroni correction are indicated in bold.

#### 3.1.3 Relationships with tissue stiffness

The relationships between volumetric strain and shear stiffness are illustrated in Figure 4, with each data point representing the subject-wise mean regional values. The corresponding regression statistics for each tissue type are detailed in Table 5. Outlier analysis excluded two regional values. Across both tissue-specific and global analyses, only weak, non-significant associations were observed between volumetric strain and shear stiffness. Subcortical and cortical GM exhibited weak negative trends, whereas white matter displayed a weak positive trend.

**Table 5.** The intercept and slope (both with 95% confidence intervals), *p-*value, and *R*^2^ value obtained from linear regressions between volumetric strain and shear stiffness using a linear mixed effects model for each tissue type.

| Region | Intercept [$10^{-4}$] | Slope [$10^{-4}$] | $p$-value | $R^2$ |
| --- | --- | --- | --- | --- |
| Global | 8.8 (3.1, 6.5) | -0.2 (-0.0, 0.4) | 0.025 | 0.22 |
| Cortical GM | 10.3 (7.7, 13.0) | -0.4 (-0.7, -0.1) | 0.016 | 0.39 |
| Subcortical GM | 13.2 (8.9, 17.5) | -0.5 (-1.3, 0.2) | 0.184 | 0.21 |
| WM | 4.8 (3.1, 6.5) | 0.2 (-0.0, 0.4) | 0.062 | 0.21 |

**Figure 4.**
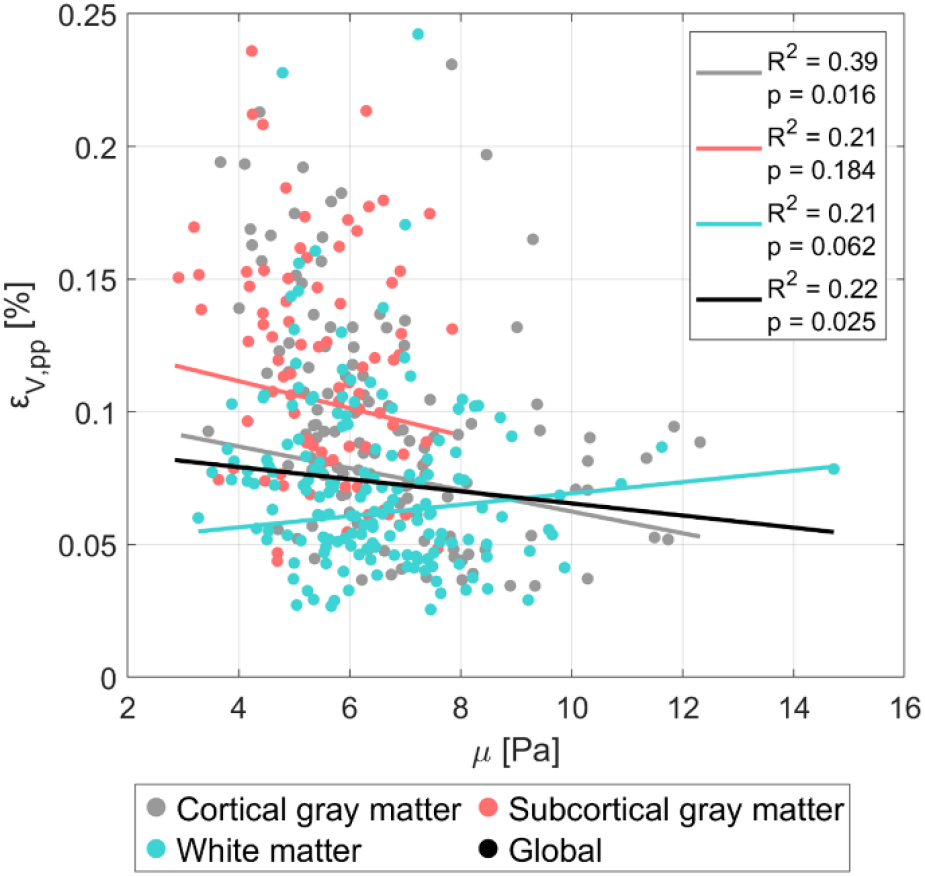
The dependence of volumetric strain on shear stiffness for cortical gray matter (gray), subcortical gray matter (red), white matter (teal). Each dot represents the subject-wise mean value across a given region, calculated only for one repeated scan. For each tissue type, the colored line shows the fixed-effect linear regression (weighted by region size) estimated from a linear mixed effects model, representing the overall population-level trend while accounting for subject- and scan-specific variability. The black line depicts the equivalent regression across all tissue types combined. The legend displays the corresponding *p*-values and *R*^2^ for each regression.

### 3.1 Octahedral shear strain

#### 3.1.1 Relationships with ATT (proxy for pulse pressure)

The relationships between octahedral strain and ATT are illustrated in Figure 5, with each data point representing the subject-wise mean regional values. The corresponding regression statistics for each tissue type are detailed in Table 6. Outlier analysis excluded one regional value. Octahedral shear strain demonstrated significant negative correlations with ATT in cortical GM, and at the global level, and a weaker negative correlation in WM. Meanwhile, a significant positive trend appeared in the subcortical GM.

**Table 6.** The intercept and slope (both with 95% confidence intervals), *p-*value, and *R*^2^ value obtained from linear regressions between octahedral shear strain and ATT using a linear mixed effects model for each tissue type. Significant correlations after Bonferroni correction are indicated in bold.

| Region | Intercept [$10^{-4}$] | Slope [$10^{-4}$] | $p$-value | $R^2$ |
| --- | --- | --- | --- | --- |
| Global | 76.5 (46.9, 61.4) | -29.5 (-11.5, -0.8) | **< 0.001** | 0.41 |
| Cortical GM | 77.6 (71.3, 84.0) | -34.2 (-39.1, -29.4) | **< 0.001** | 0.74 |
| Subcortical GM | 47.7 (39.9, 55.5) | 13.2 (6.4, 19.9) | **< 0.001** | 0.70 |
| WM | 54.2 (46.9, 61.4) | -6.2 (-11.5, -0.8) | 0.024 | 0.54 |

**Figure 5.**
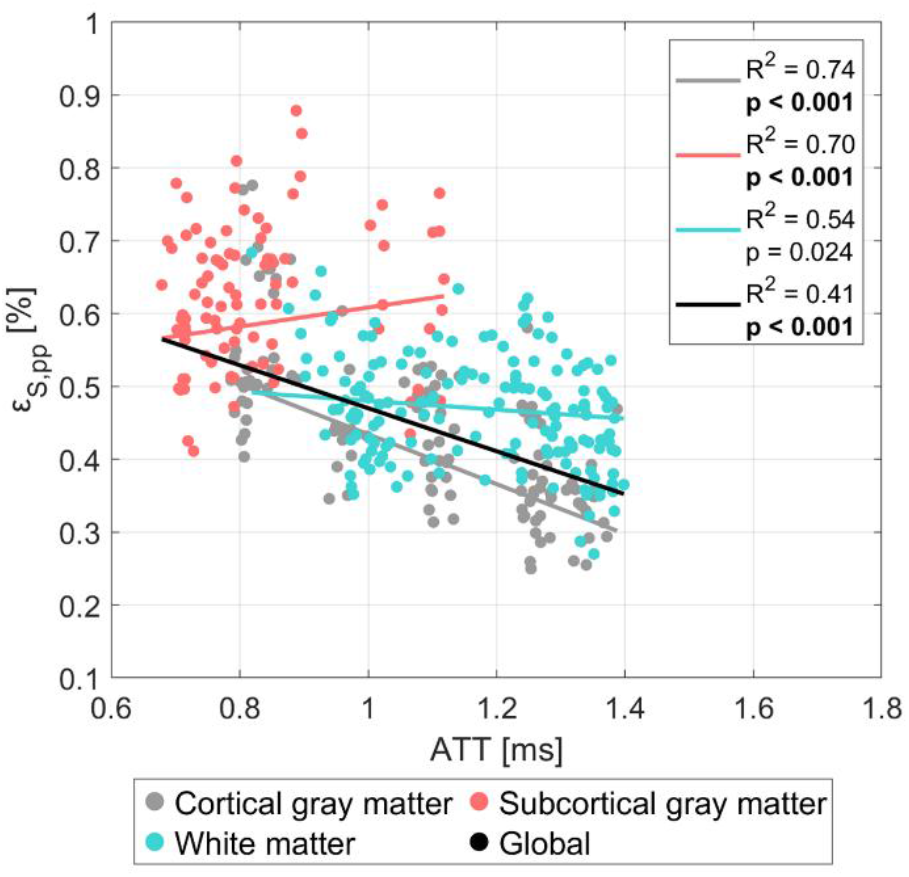
The dependence of octahedral shear strain on ATT for cortical gray matter (gray), subcortical gray matter (red), white matter (teal). Each dot represents the subject-wise mean value across a given region, calculated only for one repeated scan. For each tissue type, the colored line shows the fixed-effect linear regression (weighted by region size) estimated from a linear mixed effects model, representing the overall population-level trend while accounting for subject- and scan-specific variability. The black line depicts the equivalent regression across all tissue types combined. The legend displays the corresponding *p*-values and *R*^2^ for each regression, where significant correlations after Bonferroni correction are indicated in bold.

#### 3.1.2 Relationships with CBV (proxy for vascular structure)

The relationships between octahedral shear strain and CBV are illustrated in Figure 6, with each data point representing the subject-wise mean regional values. An outlier analysis was performed for the CBV comparison, with three regional values excluded. The corresponding regression statistics for each tissue type are detailed in Table 7. Linear regression analysis revealed a significant negative correlation between octahedral shear strain and CBV globally, but a significant positive correlation in WM. Both cortical and subcortical gray matter displayed minimal dependence on CBV.

**Table 7.** The intercept and slope (both with 95% confidence intervals), *p-*value, and *R*^2^ value obtained from linear regressions between octahedral shear strain and CBV using a linear mixed effects model for each tissue type. Significant correlations after Bonferroni correction are indicated in bold.

| Region | Intercept [$10^{-4}$] | Slope [$10^{-4}$] | $p$-value | $R^2$ |
| --- | --- | --- | --- | --- |
| Global | 51.8 (30.2, 40.5) | -4.6 (5.3, 11.3) | **< 0.001** | 0.21 |
| Cortical GM | 38.8 (28.7, 48.8) | -0.6 (-4.6, 3.4) | 0.764 | 0.24 |
| Subcortical GM | 61.9 (52.3, 71.4) | -1.2 (-4.7, 2.3) | 0.493 | 0.65 |
| WM | 35.3 (30.2, 40.5) | 8.3 (5.3, 11.3) | **< 0.001** | 0.59 |

**Figure 6.**
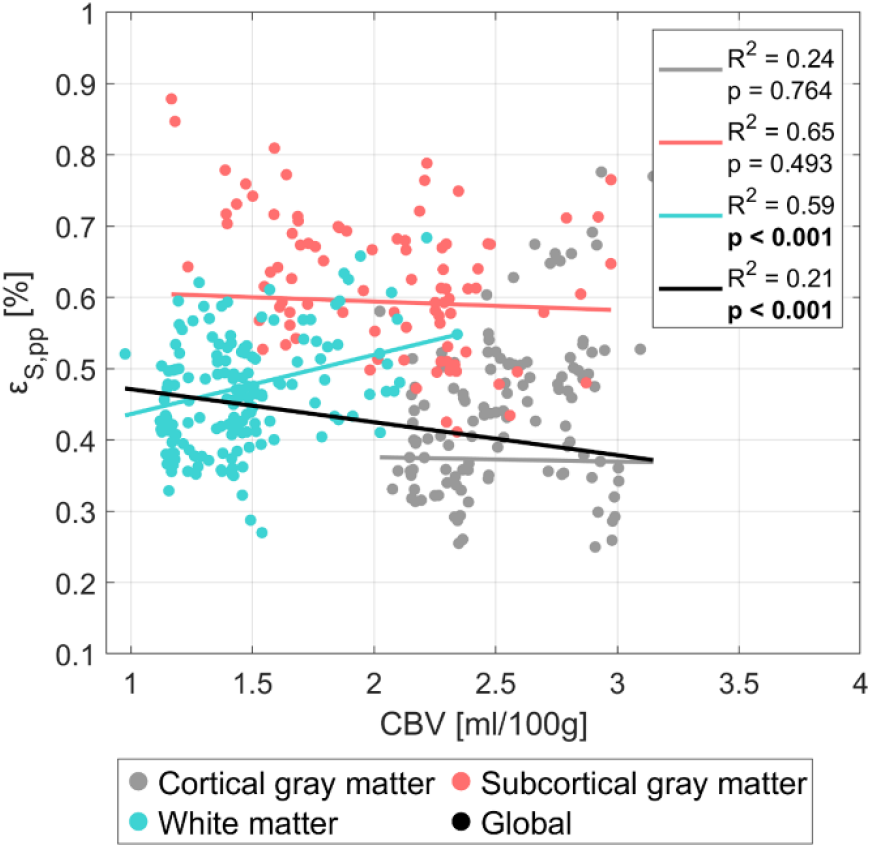
The dependence of octahedral shear strain on CBV for cortical gray matter (gray), subcortical gray matter (red) and white matter (teal). Each dot represents the subject-wise mean value across a given region, calculated only for one repeated scan. For each tissue type, the colored line shows the fixed-effect linear regression (weighted by region size) estimated from a linear mixed effects model, representing the overall population-level trend while accounting for subject- and scan-specific variability. The black line depicts the equivalent regression across all tissue types combined. The legend displays the corresponding *p*-values and *R*^2^ for each regression, where significant correlations after Bonferroni correction are indicated in bold.

#### 3.1.3 Relationships with tissue stiffness

The relationships between octahedral strain and tissue stiffness are illustrated in Figure 7, with each data point representing the subject-wise mean regional values. The corresponding regression statistics for each tissue type are detailed in Table 8. Outlier analysis excluded one regional value. Octahedral shear strain showed significant negative correlations to shear stiffness across subcortical GM, WM and globally, and a weak negative correlation in cortical gray matter.

**Table 8.** The intercept and slope (both with 95% confidence intervals), *p-*value, and *R*^2^ value obtained from linear regressions between octahedral shear strain and shear stiffness using a linear mixed effects model for each tissue type. Significant correlations after Bonferroni correction are indicated in bold.

| Region | Intercept [$10^{-4}$] | Slope [$10^{-4}$] | $p$-value | $R^2$ |
| --- | --- | --- | --- | --- |
| Global | 61.9 (52.1, 60.4) | -2.9 (-2.0, -1.0) | **< 0.001** | 0.30 |
| Cortical GM | 44.4 (37.8, 51.0) | -1.0 (-1.9, -0.1) | 0.025 | 0.25 |
| Subcortical GM | 83.1 (76.1, 90.1) | -4.3 (-5.5, -3.2) | **< 0.001** | 0.75 |
| WM | 56.2 (52.1, 60.4) | -1.5 (-2.0, -1.0) | **< 0.001** | 0.57 |

**Figure 7.**
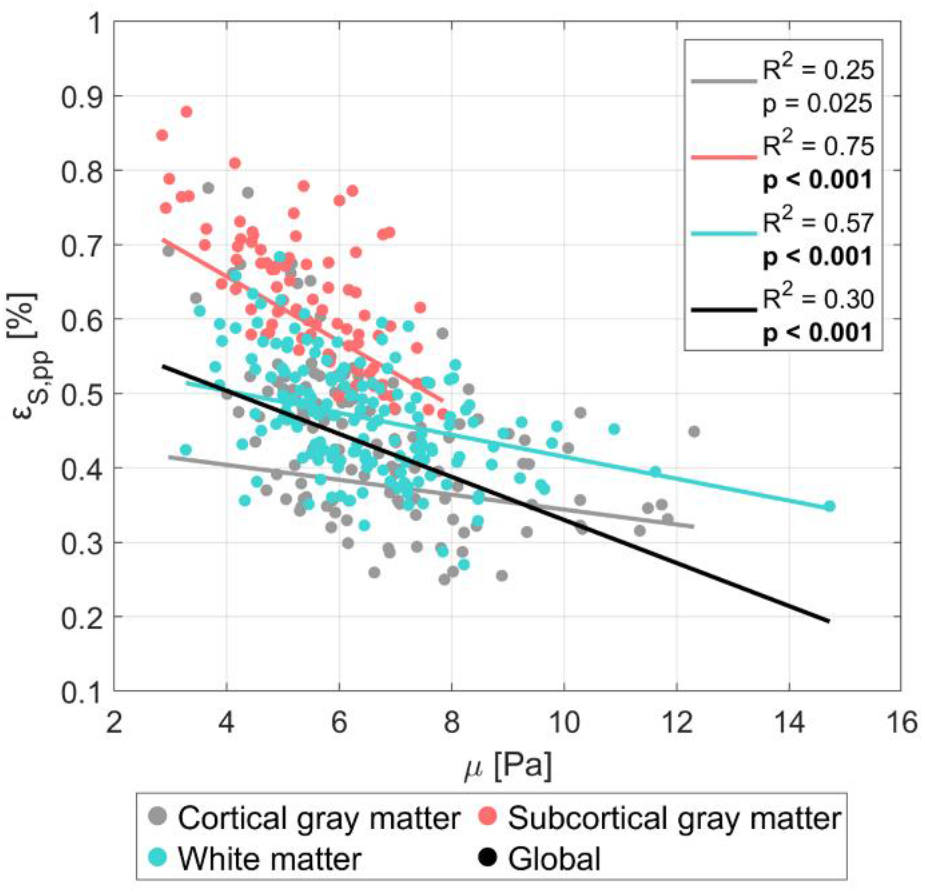
The dependence of octahedral shear strain on shear stiffness for cortical gray matter (gray), subcortical gray matter (red), white matter (teal). Each dot represents the subject-wise mean value across a given region, calculated only for one repeated scan. For each tissue type, the colored line shows the fixed-effect linear regression (weighted by region size) estimated from a linear mixed effects model, representing the overall population-level trend while accounting for subject- and scan-specific variability. The black line depicts the equivalent regression across all tissue types combined. The legend displays the corresponding *p*-values and *R*^2^ for each regression, where significant correlations after Bonferroni correction are indicated in bold.

## 4 Discussion

In this study, we combined strain tensor imaging (STI), intrinsic MR elastography (iMRE), and standard-space atlas maps in a regional analysis to explore potential relationships between CAP-related strain metrics and underlying drivers related to pulse pressure (ATT), vascular properties (CBV), or brain tissue (shear stiffness). While the use of regional correlations inherently limits the ability to draw causal conclusions, the results offer indications that contribute to a better understanding of the drivers behind the measured strains. The diversity of the results underscores the complexity of the relationship between vascular or intervascular tissue properties and the resulting measured strains. In the following section, we elaborate on these observations and consider their potential implications for the understanding of CAP-related strains in the human brain.

### 4.1 Relationships with ATT

The pulsatile pressure gradient arising from the cardiac pulse is expected to be the highest at the moment of arrival in the brain, gradually decreasing as the pulsatile energy is absorbed by the downstream vasculature (Arts et al., 2022). While the arterial transit time (ATT) is not a direct metric for pulse pressure, it serves as a meaningful biological proxy for this explorative study. Consequently, volumetric strain is anticipated to decrease with increasing ATT, a trend observed in both cortical GM, WM, and globally. This highlights the relationship between the pulse pressure and volumetric strain and partly explains why volumetric strain is higher in GM (which is perfused earlier than WM) compared to WM. A similar relationship was found in octahedral shear strain, indicating that the pulse pressure is an important component in both strain metrics and serves as a common driver. However, this pattern was not observed in subcortical GM, where volumetric strain showed no clear dependence on ATT and octahedral shear strain exhibited the opposite trend. This can partly be explained by subcortical GM containing few regions where the cardiac pulse reaches each region at similar arrival times. This likely reduces the ability to detect clear relationships between ATT and strain metrics in this region, consequently making this region more sensitive to noise. However, it could also hint at more complex behavior arising from, for example, unique vascular architecture and/or mechanical coupling with surrounding tissues.

### 4.2 Relationships with vasculature

Previous studies have suggested that CBV may be a key factor in explaining the differences in volumetric strain between gray and white matter, given that similar disparities are observed in CBV (Adams et al., 2019). While our results show a modest correlation between volumetric strain and CBV, the CBV maps used in this analysis include the combined contributions of arterial, venous, and capillary blood volumes. Since most of the pulsatility in the vasculature is driven by arterial vessels (which only comprise 20-30% of the total volume (Hua et al., 2019)), with only minimal pulsations occurring in capillaries, a map that isolates arterial CBV would potentially provide a more accurate explanation of the observed volumetric strain differences. Nonetheless, it is unlikely that the blood volume alone can explain the observed 2.3 factor difference in volumetric strain between gray and white matter. In contrast, octahedral shear strain showed considerably less dependence on CBV.

### 4.3 Relationships with tissue stiffness

Differences in gray and white matter tissue stiffness have previously been suggested as a potential explanation for disparities in volumetric strain (Adams et al., 2019). However, our results indicate little to no correlation between shear stiffness and volumetric strain across tissue types, suggesting that tissue stiffness contributes minimally to volumetric strain. In contrast, octahedral shear strain demonstrated a significant negative correlation with shear stiffness in both subcortical gray matter and white matter, consistent with the expected inverse relationship between shear strain and material stiffness. Furthermore, octahedral shear strain showed considerably less overlap across tissue types when plotted against the properties of interest compared to volumetric strain. As each tissue type is expected to exhibit distinct tissue properties, this indicates that octahedral shear strain has a stronger dependence on such tissue properties. Consistent with this interpretation, linear regressions for octahedral shear strain against stiffness yielded generally higher *R*^2^ values when performed separately within each tissue type, emphasizing the appropriateness of this stratification for stiffness-related effects. In comparison, volumetric strain showed less improvement with tissue-specific stratification, consistent with the idea that vascular metrics play a larger role in volumetric strain, which may vary less distinctly across tissue types.

### 4.4 Interpretation of strain metrics

Although it is known that cerebral arterial pulsations induce the measured strain in the brain, the interpretation of the observed strain metrics is still uncertain. Previous work suggested that volumetric strain reflects the swelling of the microvasculature embedded in the tissue (Adams et al., 2019), placing volumetric strain as a potential important metric to assess the state of the microvasculature. Our results indicate volumetric strain being associated with mainly CBV and the pulse pressure, while showing minimal dependence on tissue stiffness. At the same time, octahedral shear strain showed less dependence on CBV, and stronger associations with the pulse pressure and tissue stiffness. While both metrics seem largely influenced by the pulse pressure, these results indicate that volumetric strain primarily reflects vascular effects while octahedral shear strain is more related to the surrounding tissue. This interpretation is further supported by a recent case study (J.-J. Sloots et al., 2023), in which STI was performed on a patient who underwent craniotomy following traumatic brain injury. In this case, the global boundary conditions of the brain were heavily influenced, which in turn greatly affected the octahedral shear strain. In contrast, volumetric strain closely resembled that of healthy volunteers, suggesting it is predominantly determined by the local properties of the vessel walls.

### 4.5 Considerations around atlas maps

We compared our values to standardized MNI atlases for certain metrics rather than direct measurements from the same subjects, which introduces a layer of potential error. While direct measurements from each subject would be preferable, the use of atlases were considered more appropriate and resource efficient for an explorative study such as this one. Nevertheless, the limited availability of atlas maps introduced variability in subject demographics. While the sex ratio and age distribution were similar for the CBF, MTT, and ATT maps, the CBV map included roughly twice as many males as females and an older overall population. Moreover, the CBV atlas was constructed from patients with grade IV glioblastoma; although tumor regions were removed from each map, it is possible that the overall CBV distribution may still be subtly affected.

### 4.6 Limitations

Both volumetric and octahedral shear strain are inherently noisy measurements, as they result from spatial derivative operations. While assessing peak-to-peak values over a certain region proved generally effective in handling noise, there is also information lost in such a simplification. In addition, a considerable number of voxels were excluded due to region erosion, artifact removal, and masking of problematic fluid flows. Thus, while the analysis covered most of the brain, it still represents only a subset of total brain volume. Additionally, we performed a regional comparison, where each region likely contains a unique proportion of underlying contributions in the measured strains. This makes it more difficult to identify direct dependencies because regional variations in tissue composition, vascular density, or microstructural organization may independently influence both strain and the other measured properties. Such co-varying regional characteristics could either produce spurious correlations where no causal link exists or, conversely, mask genuine associations, resulting in the generally low *R*^2^ values observed in the linear regressions. Finally, because the stiffness and strain metrics were derived from the same underlying displacement data, there remains a potential risk of spurious correlations arising from shared data dependencies. Despite this, the substantially different methods employed to estimate stiffness and strain metrics appeared to be robust against such correlations, as indicated by the lack of significant association between shear stiffness and volumetric strain.

### 4.7 Brain strain is driven by complex interplay

While our results help to shed light on possible drivers of volumetric and octahedral shear strain, they also underscore the complexity of these metrics. The observed associations suggest that both strain metrics reflect an intricate interplay of multiple physiological and structural factors, rather than being directly governed by any single metric. In particular, while CBV shows a notable relationship with volumetric strain, this likely represents only part of a broader system of influences. Not only do vascular wall compliance and pulse pressure seem to play important roles, but we further speculate that vascular architecture, such as branching patterns or hydraulic permeability (i.e., the ease with which fluid can move through the vascular network, as described in poroelasticity), also influences strain dynamics. As hydraulic permeability decreases, vascular resistance increases, potentially leading to greater energy dissipation, much like how electrical resistance in a circuit converts energy into heat. This dissipation may, in turn, modulate strain behavior in the brain. Given this complexity, a more comprehensive characterization of the underlying principles may require a system modeling approach, which could integrate the influences of various mechanical, vascular, and boundary condition effects to provide a more complete understanding of brain tissue deformation dynamics. Future research is needed to further unravel these interactions and their implications for brain biomechanics in both health and disease.

## 5 Conclusion

This study provides preliminary insights into the potential processes that may contribute to volumetric and octahedral shear strain in the human brain, suggesting that these strain metrics could reflect different physiological processes. Volumetric strain showed stronger associations with cerebral blood volume, while octahedral shear strain appeared more closely related to tissue stiffness. However, both strain metrics appear to arise from a more intricate interplay of factors than previously assumed. Within the context of a large, complex organ such as the brain, these effects underscore the need for further research to fully characterize the dynamics of brain strain. These findings lay the groundwork and can provide benchmarks for e.g. computational modeling studies to further investigate of the physiological determinants of strain metrics.

## Acknowledgements

The authors would like to thank Dr. Alex A. Bhogal and Dr. Frederick B. Blaise for guidance with the atlas maps used in this study.

