## Supplementary Material for "Exploring the role of vascular factors and tissue properties in pulsatile brain deformation"

### Supplementary Files

#### ATLAS overview

**Table S1:** Overview of sources and subject characteristics for each atlas map used in the analysis.

| Metric | Reference | Source | Number of subjects | Sex (F/M) | Age | Health status |
|---|---|---|---|---|---|---|
| CBV | Fuster-Garcia et al., 2021 | NCT03439332 clinical study | 134 | 66/118* | 24 – 81 (mean age: 60) years* | Glioblastoma grade IV** |
| MTT | Frederick B. Blaise, 2016 | HCP1200 | 1206 | 656/550 | 22 – 35 years | Healthy subjects |
| CBF, ATT | Taso et al., 2021 | Measured in Taso et al., 2021 | 10 | 6/4 | 30 ± 7 years | Healthy subjects |

*A subset of 134 subjects was used to generate the final CBV map. Sex and age information were available only for the initial 184 subjects.

**Tumor regions were excluded from each subject's map.

#### Relationship between volumetric strain and hemodynamic metrics

The relationships between volumetric strain and the hemodynamic properties of interest are illustrated in Figure S1, with each data point representing the subject-wise mean regional values. The corresponding regression statistics for each tissue type and property are detailed in Table S2. A separate outlier analysis was performed for each volumetric strain comparison. Three regional values were excluded for the regressions with CBF, and two for the MTT. Linear regression analysis revealed significant positive correlations between volumetric strain and CBF in white matter and globally. Meanwhile, weak negative trends were observed in cortical and subcortical GM, potentially highlighting distinct tissue-specific dependencies. A significant negative correlation was found for MTT on the global level while a significant positive trend was observed for the subcortical GM.

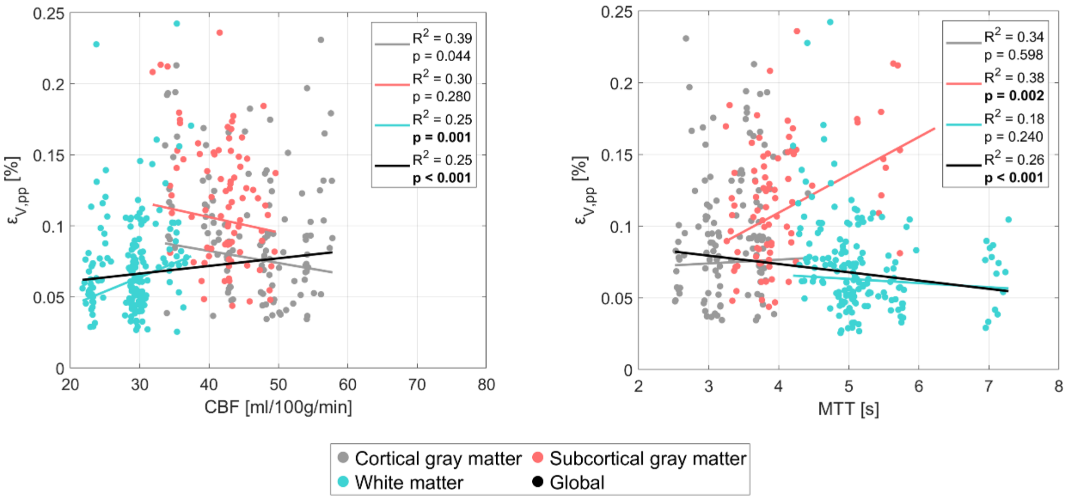


**Figure S1:** The dependence of volumetric strain on hemodynamic properties for cortical gray matter (gray), subcortical gray matter (red), white matter (teal). Each dot represents the subject-wise mean value across a given region, calculated only for one repeated scan. For each tissue type, the colored line shows the fixed-effect linear regression (weighted by region size) estimated from a linear mixed effects model, representing the overall population-level trend while accounting for subject- and scan-specific variability. The black line depicts the equivalent regression across all tissue types combined. The legend displays the corresponding $p$-values and $R^2$ for each regression, where significant correlations after Bonferroni correction are indicated in bold.

**Table S2:** The intercept and slope (both with 95% confidence intervals), $p$-value, and $R^2$ value obtained from linear regressions between volumetric strain and the hemodynamic properties of interest using a linear mixed effects model for each tissue type. Significant correlations after Bonferroni correction are indicated in bold.

| | Region | Intercept $[10^{-4}]$ | Slope $[10^{-4}]$ | $p$-value | $R^2$ |
|---|---|---|---|---|---|
| *CBF* | Global | 5.0 (-3.2, 3.9) | 0.1 (0.1, 0.3) | **< 0.001** | 0.25 |
| | Cortical GM | 11.6 (7.4, 15.9) | -0.1 (-0.2, -0.0) | 0.044 | 0.39 |
| | Subcortical GM | 14.9 (6.5, 23.4) | -0.1 (-0.3, 0.1) | 0.280 | 0.30 |
| | WM | 0.4 (-3.2, 3.9) | 0.2 (0.1, 0.3) | **0.001** | 0.25 |
| *MTT* | Global | 9.7 (5.0, 10.5) | -0.6 (-0.8, 0.2) | **< 0.001** | 0.26 |
| | Cortical GM | 6.6 (3.1, 10.1) | 0.3 (-0.7, 1.3) | 0.598 | 0.34 |
| | Subcortical GM | 0.4 (-6.1, 6.9) | 2.6 (1.0, 4.3) | **0.002** | 0.38 |
| | WM | 7.8 (5.0, 10.5) | -0.3 (-0.8, 0.2) | 0.240 | 0.18 |

#### Relationship between octahedral shear strain and hemodynamic metrics

The relationships between octahedral strain and the hemodynamic properties of interest are illustrated in Figure S2, with each data point representing the subject-wise mean regional values. An outlier analysis was performed separately for each octahedral shear strain comparison. Two regional values were excluded from the comparisons with CBF and MTT. The corresponding regression statistics for each tissue type and property are detailed in Table S3. CBF showed significant positive correlations in WM, while a significant negative correlation was identified at the global level. For MTT, significant positive trends were observed both globally and within cortical GM, while significant negative correlation was observed in WM. Both CBF and MTT exhibited markedly different behaviors across tissue types, with minimal overlap among data points. In general, octahedral shear strain regressions yielded considerably higher R^2 values compared to volumetric strain regressions for hemodynamic comparisons.

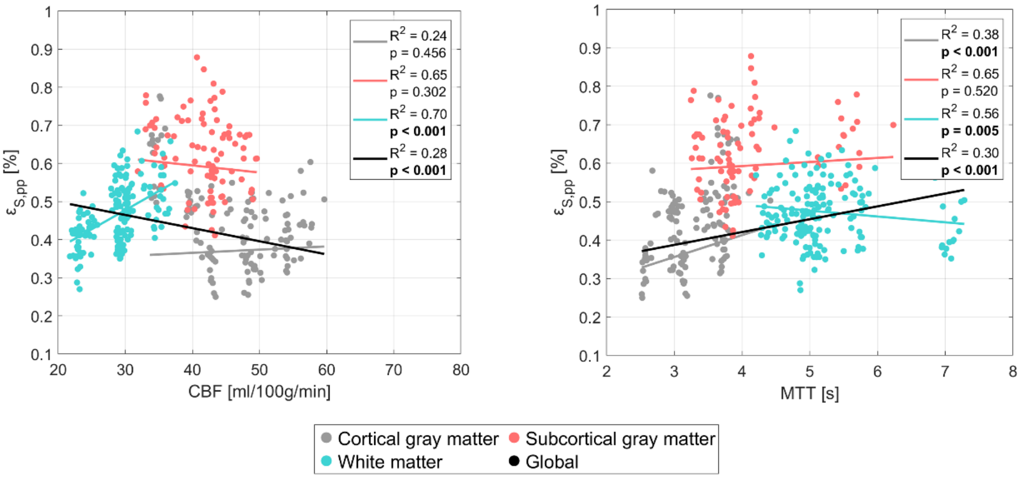


**Figure S2:** The dependence of octahedral shear strain on hemodynamic properties for cortical gray matter (gray), subcortical gray matter (red) and white matter (teal). Each dot represents the subject-wise mean value across a given region, calculated only for one repeated scan. For each tissue type, the colored line shows the fixed-effect linear regression (weighted by region size) estimated from a linear mixed effects model, representing the overall population-level trend while accounting for subject- and scan-specific variability. The black line depicts the equivalent regression across all tissue types combined. The legend displays the corresponding $p$-values and $R^2$ for each regression, where significant correlations after Bonferroni correction are indicated in bold.

**Table S3:** The intercept and slope (both with 95% confidence intervals), $p$-value, and $R^2$ value obtained from linear regressions between octahedral shear strain and the hemodynamic properties of interest using a linear mixed effects model for each tissue type. Significant correlations after Bonferroni correction are indicated in bold.

| | Region | Intercept $[10^{-4}]$ | Slope $[10^{-4}]$ | $p$-value | $R^2$ |
|---|---|---|---|---|---|
| *CBF* | Global | 56.8 (10.5, 24.9) | -0.3 (0.8, 1.2) | **< 0.001** | 0.28 |
| | Cortical GM | 33.2 (21.8, 44.5) | 0.1 (-0.1, 0.3) | 0.456 | 0.24 |
| | Subcortical GM | 67.0 (51.0, 83.0) | -0.2 (-0.5, 0.2) | 0.302 | 0.65 |
| | WM | 17.7 (10.5, 24.9) | 1.0 (0.8, 1.2) | **< 0.001** | 0.70 |
| *MTT* | Global | 28.7 (48.5, 61.9) | 3.3 (-2.6, -0.5) | **< 0.001** | 0.30 |
| | Cortical GM | 18.5 (10.3, 26.8) | 5.7 (3.3, 8.0) | **< 0.001** | 0.38 |
| | Subcortical GM | 55.1 (41.9, 68.3) | 1.1 (-2.2, 4.3) | 0.520 | 0.65 |
| | WM | 55.2 (48.5, 61.9) | -1.5 (-2.6, -0.5) | **0.005** | 0.56 |
